# Lack of food intake during shift work alters the heart transcriptome and leads to cardiac fibrosis and inflammation in rats

**DOI:** 10.1101/2021.05.13.444001

**Authors:** Alexandra J. Trott, Ben J Greenwell, Tejas R. Karhadkar, Natali N. Guerrero-Vargas, Carolina Escobar, Ruud M Buijs, Jerome S Menet

**Affiliations:** Department of Biology, Texas A&M University, College Station, TX 77843 USA; Program of Genetics, Texas A&M University, College Station, TX 77843, USA; Center for Biological Clock Research, Texas A&M University, College Station, TX 77843, USA; Departamento de Anatomía, Facultad de Medicina, Universidad Nacional Autónoma de México, Ciudad Universitaria, Mexico City, Mexico; Instituto de Investigaciones Biomédicas, Universidad Nacional Autónoma de México, Ciudad Universitaria, Mexico City, Mexico

**Keywords:** shift work, circadian rhythms, cardiovascular diseases, heart, transcriptome, RNA-Seq, cardiac fibrosis, inflammation

## Abstract

Many epidemiological studies revealed that shift work is associated with increased risk of cardiovascular diseases. However, the underlying mechanisms remain poorly understood. An experimental model of shift work in rats has been shown to recapitulate the metabolic disorders observed in human shift workers, and used to demonstrate that restricting food consumption outside working hours prevents shift work-associated obesity and metabolic disturbance. Here we used this model to characterize the effects of shift work in the heart. We show that experimental shift work reprograms the heart cycling transcriptome independently of food consumption. While phases of rhythmic gene expression are distributed across the 24-hour day in control rats, they are clustered towards discrete times in shift workers. Additionally, preventing food intake during shift work affects the expression level of hundreds of genes in the heart. Many of them are found in transcriptional signatures associated with pressure overload and cardiac hypertrophy, and encode for components of the extracellular matrix and inflammatory markers. Consistent with this, the heart of shift worker rats not eating during work exhibits fibrosis and is colonized by immune cells. While maintaining food access during shift work has less effects on gene expression, genes found in transcriptional signatures of cardiac hypertrophy remain affected, and the heart of shift worker rats exhibits fibrosis without inflammation. Together, our findings provide insights into how shift work affects cardiac function, and suggest that some interventions aiming at mitigating metabolic disorders in shift workers may have adverse effects on cardiovascular diseases.

## Introduction

Shift work is an employment practice designed to provide services outside the daytime working hours, including evening or night shifts, early morning shifts, and rotating shifts. Shift work is becoming increasingly popular in modern society, mostly because of the higher demand for productivity (1). About 15-20% of the workforce is engaged in some kind of shift work worldwide, and this number is rapidly increasing (1). Over the last decades, many epidemiological studies revealed that shift work affects human health and increases the risk of developing cancer, metabolic disorders, cardiovascular diseases, and mood disorders (1-14). Relevant to this paper, a meta-analysis performed on 17 studies concluded that shift workers exhibit a 40% increased risk of developing cardiovascular diseases than day workers (15). A more recent meta-analysis on 173,000 individuals similarly found that shift/night workers have an increased risk of cardiovascular diseases than day workers (13). However, there was significant heterogeneity between studies considered in these meta-analyses, and it is still unclear to which extent the quantity, the duration, and the type of shift work impair cardiovascular functions (10, 12).

While the mechanisms by which shift work causes cardiovascular diseases remain unclear, they likely involve a misalignment between internal biological rhythms and the natural light:dark cycle (16). In mammals, every tissue harbors a timekeeping mechanism, the circadian clock, that drives rhythms in gene expression to allow biological functions to perform optimally at the most appropriate time of the day. Circadian clocks in peripheral tissues are synchronized to the daily environmental variation by the master circadian pacemaker located in the suprachiasmatic nucleus (SCN) of the hypothalamus, which is itself entrained to the light:dark cycle via direct retinal innervation (17). The SCN utilizes multiple cues to synchronize peripheral clocks, including rhythms in neuronal signaling, hormone secretion, body temperature, and food intake. Alteration in the timing of these cues by shift work disrupts the proper functioning of peripheral clocks and clock-controlled biological functions. Accumulating evidence indicates that alteration of the daily rhythm of food intake, which is prevalent in shift workers, is particularly associated with circadian rhythm disorders (18, 19). For example, rats subjected to shift work paradigms for five weeks shift their food intake to the work phase (*i.e*., natural rest phase), and exhibit increased fat content and impaired glucose tolerance (20, 21). Importantly, these effects are not observed if rats cannot eat during work, indicating that the adverse effects of shift work on metabolic functions are mostly mediated by a shift in the timing of food intake to the natural rest phase (20, 21). Whether rats subjected to shift work also develop cardiovascular diseases, and whether those can be prevented by restricting food intake during the normal active phase, remain however unexplored.

To get insights into how shift work alters heart biology, we used a model of shift work in rats that mimics shift work in humans (22). Rats were housed in a slowly rotating wheel for 8 hours during their sleeping phase from Monday to Friday, and left undisturbed outside their work schedule. Because restricting food intake to the normal active phase prevents the development of metabolic disorders (20, 21), we hypothesized that preventing access to food during the daily 8 hours of shift work might also hinder alterations of cardiac functions. Our results indicate that five weeks of shift work in rats leads to cardiac fibrosis, and that the adverse effects of shift work in the heart are surprisingly exacerbated when shift worker rats do not have access to food during work. Our findings thus suggest that lack of food intake during shift work, an intervention that prevents metabolic disorders in shift workers, has adverse effects on cardiovascular functions.

## Results

To characterize the effects of shift work in the heart, we used a rat model of shift work previously shown to cause metabolic disorders similar to those observed in human shift workers (20, 22). Because restricting food access to the natural active phase prevents the development of obesity and diabetes in this model, we also aimed to investigate the effects of lack of food intake during work. (Fig. 1) (20). A total of 54 rats maintained on a 12:12 light:dark cycle (LD12:12, light on from 7 am to 7 pm) were randomly assigned to control (C), shift work (W), or shift work with restricted nighttime feeding (WRF) groups prior the start of the experiment. Shift work consisted of rats being placed in a slowly rotating drum (1 revolution per 3 minutes) from 9 am to 5 pm, Monday to Friday, for a total of 5 weeks (Fig. 1), as published previously (20, 22). Control rats had access to food and water *ad libitum* throughout the experiment. Worker rats had access to *ad libitum* food and water and were exposed to shift work for 5 weeks. WRF rats were also exposed to shift work and had *ad libitum* access to water, but only had access to food when not doing shift work (*i.e*., from 5 pm to 9 am; Fig. 1). For each group, rats were left undisturbed and had free access to food and water during weekends. After 5 weeks of shift work, rats were euthanized on a Saturday under a weekend schedule, and the hearts collected and flash-frozen.

**Fig. 1.**
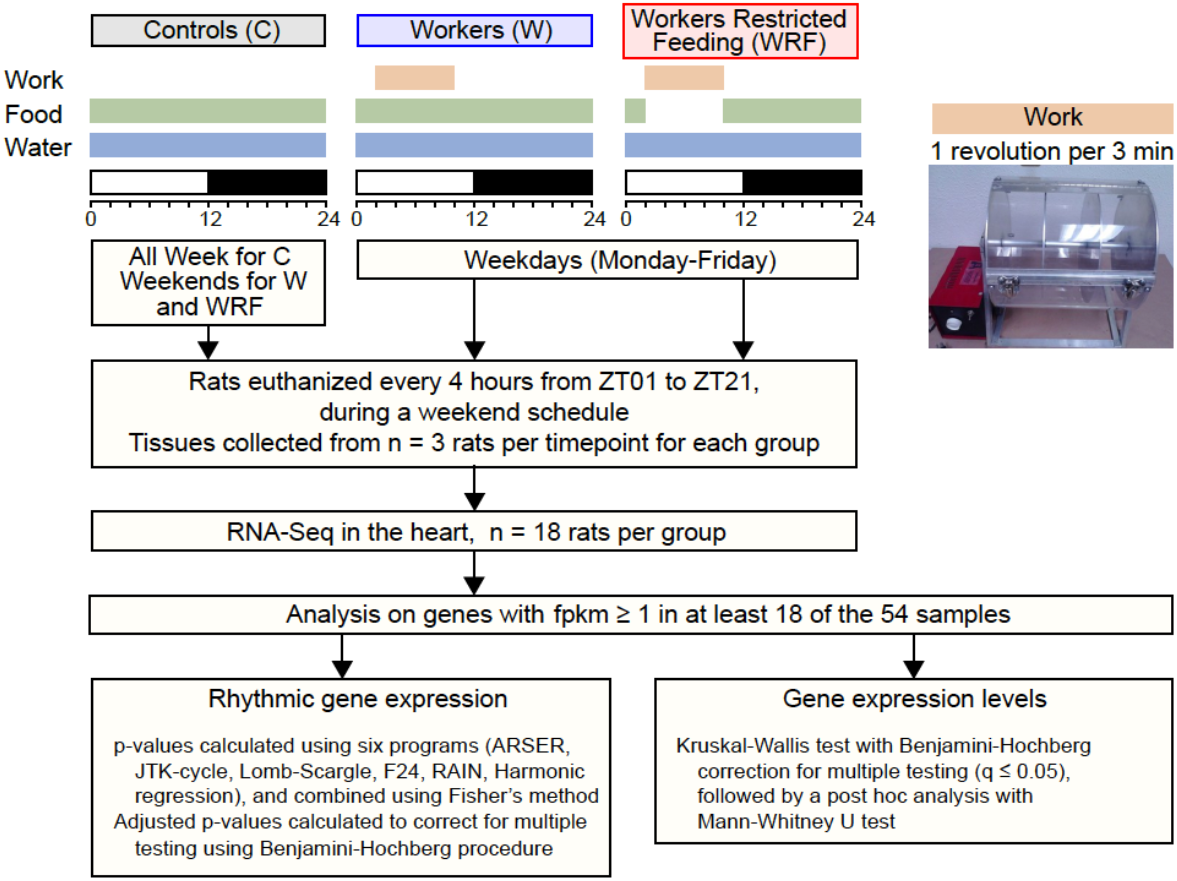
Experimental setup. Individually housed rats exposed to LD12:12 were randomly assigned to one of the three groups: control (C), shift worker (W), or shift worker subjected to restricted feeding during work (WRF). All rats had ad libitum access to water. C rats had ad libitum access to food and were not manipulated throughout the experiment. W and WRF rats were placed in a rotating drum (1 revolution per 3 minutes) for eight hours (9 am-5 pm, *i.e*., ZT2 to ZT10), Monday-to-Friday, for 5 consecutive weeks to simulate shift work, and were maintained unrestrained in their cages over the weekend. W rats had *ad libitum* access to food at all times, including during work, whereas WRF rats only had access to food outside the work schedule, *i.e*., from 5 pm to 9 am. After five weeks, rats were euthanized during a week-end schedule (no work) every four hours with n=3 rats per timepoint and group. Hearts were collected, flash-frozen, and processed afterward for RNA extraction and RNA-Seq library preparation.

### Shift work reprograms the heart cycling transcriptome

To determine the effects of shift work in the heart, we first performed an RNA-Seq analysis of the heart transcriptome across the 24-hour day in C, W, and WRF rats (6 timepoints, n = 3 biological replicates per timepoint and group). Analysis of rhythmic gene expression revealed large differences in the number of rhythmically expressed genes, with the number of rhythmic genes being ∼2-fold higher in WRF rats than in C rats, and ∼2-fold lower in W rats than in C rats (Fig. 2A, Table S1). Rhythmically expressed genes poorly overlapped between the three groups, suggesting that shift work profoundly reprograms the rat heart cycling transcriptome (Fig. 2A, 2B). Interestingly, while phases of rhythmic gene expression in C rats were distributed across the 24-hour day with most genes peaking between ZT0 and ZT20, rhythmic genes in both W and WRF rats displayed discrete peak phases of expression, *i.e*., ZT11-14 for W rats, and ZT7-9 and ZT20-21 in WRF rats (Fig. 2B, 2C). This suggests that shift work, which occurs recurrently from ZT2 to ZT10, stimulates gene expression at specific phases of the day, and that this peak of expression persists the day following shift work when animals are left undisturbed, *i.e*., when heart samples were collected on a Saturday (Fig. 2A).

**Fig. 2.**
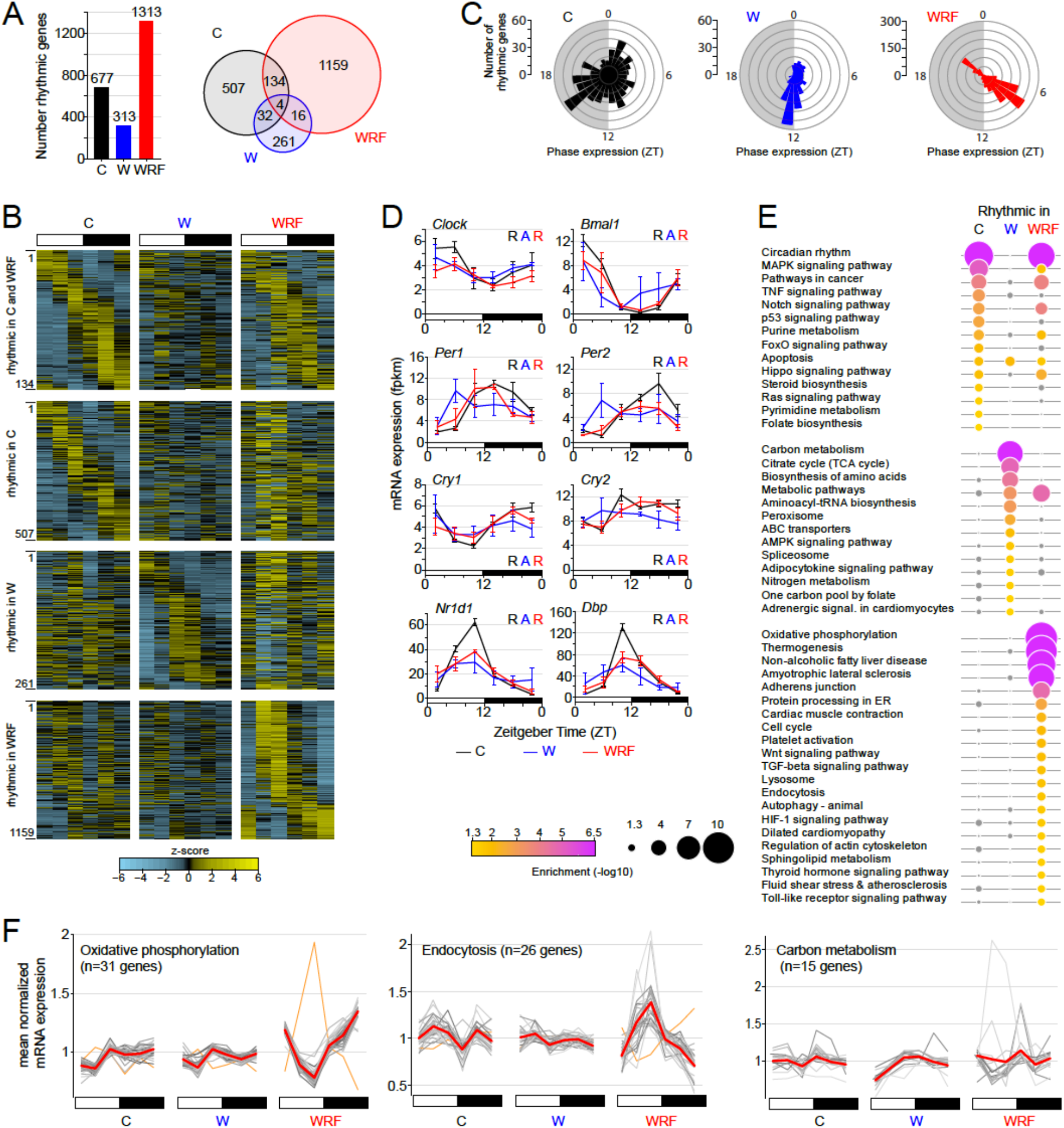
Effect of shift work on the cycling heart transcriptome. **A**. Total number (left) and overlap (right) between rhythmically expressed genes for each group: control (C, black), shift worker (W, blue), or shift worker subjected to restricted feeding during work (WRF, red). **B**. Heatmap visualization of rhythmic gene expression for genes rhythmic in both C and WRF groups, C only, W only, and WRF only. The signal for each timepoint corresponds to the average of 3 individual samples. Genes were ordered based on the phase of rhythmic gene expression of C rats for the top two heatmaps, while they were ordered based on the phase of rhythmic gene expression in W and WRF groups for genes rhythmic in W only and WRF only, respectively. **C**. Rose plot representation of the phase of rhythmic gene expression in C, W, and WRF rat groups. **D**. Clock gene rhythmic expression in the heart of C, W, and WRF rats. The signal corresponds to the mean +/- s.e.m of 3 animals per timepoint. Statistical analysis for rhythmic gene expression is displayed as R (rhythmic expression) or A (arrhythmic expression), and the font color refers to each group: C, black; W, blue; WRF, red. **E**. KEGG pathway enrichment for genes being rhythmically expressed in C, W, and WRF rat heart. Pathways that are not enriched in certain groups (p > 0.05) are displayed in grey. **F**. Expression profile of genes accounting for the enrichment of three KEGG pathways. The expression of individual genes is displayed in grey, and the averaged expression in red. For the oxidative phosphorylation and endocytosis pathways (WRF enriched pathways), the rhythmic expression of one gene is in antiphase to all other genes; this gene is displayed in orange and was not taken into account for calculating the averaged expression.

Examination of core clock gene expression showed that shift work alone impaired the molecular clock oscillation in the heart, and that preventing food access during shift work restored rhythmic gene expression (Fig. 2D). However, the amplitude of rhythmic expression in WRF rats was significantly decreased for most clock genes (Fig. 2D, S1A). Moreover, arrhythmic expression of clock genes in W heart was mostly due to differences in peak expression between biological replicates, with the replicate rhythm 2 showing out-of-phase rhythmic expression compared to the replicate rhythms 1 and 3 (Fig. S1B). While it remains unclear why the molecular clock oscillation is shifted in only one replicate rhythm, it is possible that the heart molecular clock is less sensitive to shift work synchronizing cues (*e.g*., increased daytime food intake) than other tissues like the liver (see discussion) (20, 22).

To determine if shift work reprograms the rhythmicity of specific biological pathways in the rat heart, we performed a KEGG pathway analysis (23) of genes being rhythmically expressed in either C, W, or WRF rats. Consistent with our finding that rhythmic genes poorly overlap between groups, most pathways were enriched in a group-specific manner (Fig. 2E, Table S2). Genes being rhythmically expressed in C rats were enriched for various signaling pathways, including MAPK, TNF, p53, and FoxO signaling pathways (Fig. 2E). Although rhythmic genes in W and WRF rats were both enriched in metabolic pathways, the nature of these pathways was group-specific, likely reflecting differences in the timing of food intake between W and WRF rats (20). Rhythmic genes in W rats were enriched in pathways involved in carbon metabolism, biosynthesis of amino acid and aminoacyl tRNA, and nitrogen metabolism, whereas rhythmic genes in WRF group were mostly involved in oxidative phosphorylation (Fig. 2E). Genes being rhythmically expressed in WRF rats were also enriched in pathways associated with membrane-enclosed organelles (*e.g*., protein processing in the endoplasmic reticulum, lysosome, endocytosis, and autophagy), as well as pathways associated with cardiac functions such as cardiac muscle contraction and dilated cardiopathy (Fig. 2E).

Given that most rhythmic genes in W and WRF rats peaked at discrete phases (Fig. 2C), we examined the expression of rhythmic genes from several enriched KEGG pathways (Fig. 2F). Remarkably, rhythmic genes from the same enriched pathways displayed similar patterns of expression (Fig. 2F, S2, Table S3). For example, the expression of 30 out of 31 rhythmic genes involved in oxidative phosphorylation was highly coordinated in WRF rats, with trough expression at ZT9 at the end of shift work. Similarly, 25 out of 26 rhythmic genes involved in endocytosis displayed similar expression profiles in WRF rats, with peak expression at the end of shift work. Moreover, 15 rhythmic genes in W rats involved in carbon metabolism showed similar profiles of expression and peaking a few hours after the end of shift work.

Taken together, our data indicate that both shift work and shift work without food intake profoundly reprogram the heart cycling transcriptome. Many genes normally expressed rhythmically in rat heart across the 24-hour day lose rhythmicity. Conversely, both shift work protocols lead to the emergence of new rhythms of gene expression, which for the most part occur at discrete phase across the day and are likely to be directly induced by the lack of food intake during shift work or by shift work alone.

### Gene expression in the heart is profoundly altered in shift workers subjected to restricted feeding

To further investigate the effects of shift work in the heart, we also performed an analysis of differential gene expression between the 3 groups (Fig. 1). This analysis revealed that 944 genes (9.2% of all expressed genes) were affected by shift work (Fig. 3A, Fig. S3, Table S4). Remarkably, the misregulation of heart gene expression by shift work was almost exclusively restricted to the WRF group (n = 904 genes; W, n = 40 genes) (Fig. 3A). In particular, the expression of 808 genes were significantly increased in WRF rats when compared to C and W rats, while only 67 genes exhibited a decreased expression (Fig. 3B).

**Fig. 3.**
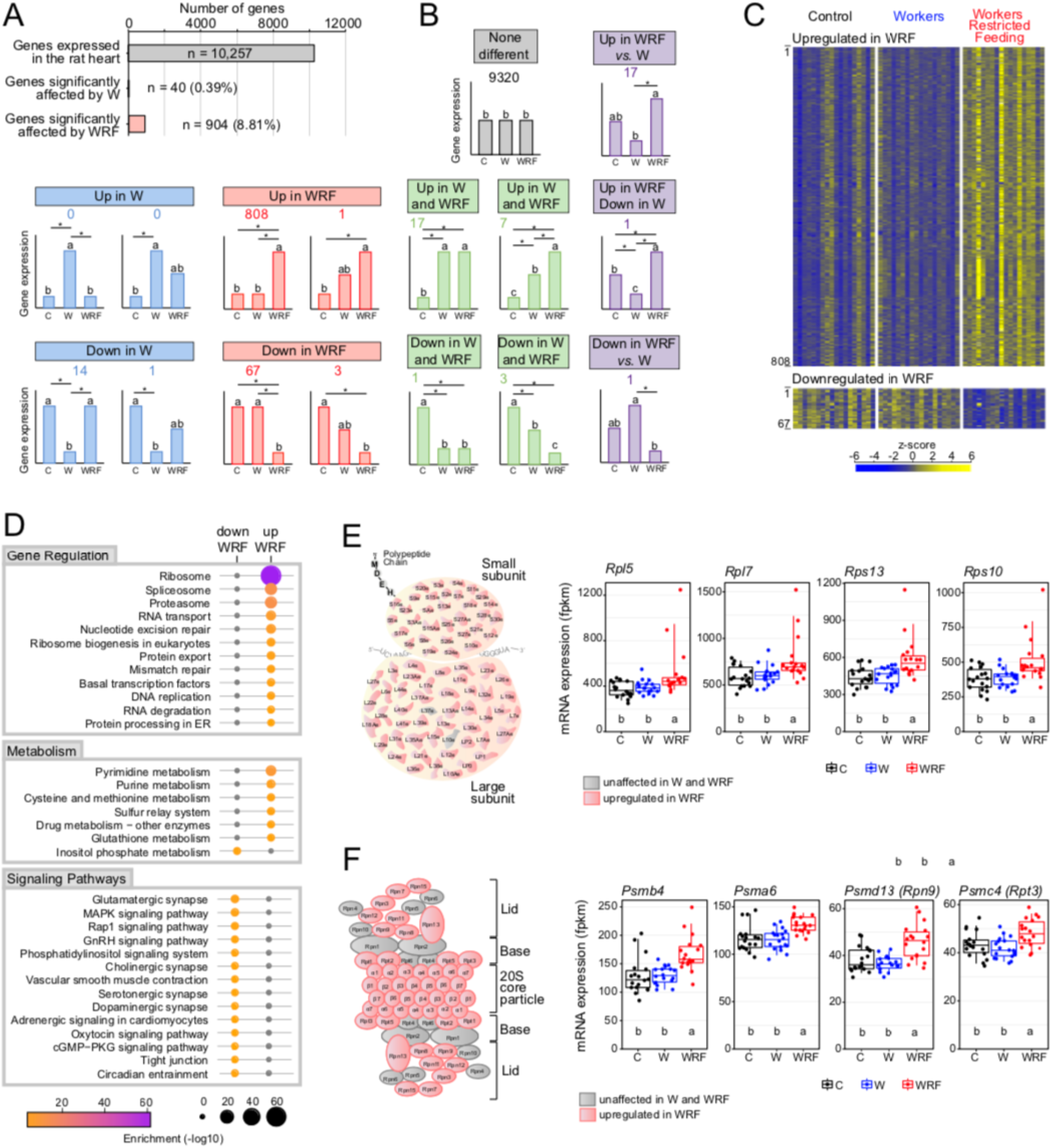
Effect of shift work on cardiac gene expression. **A, B**. Number of differentially expressed genes between C, W, and WRF groups (A), and description of the significant effects between groups after Mann-Whitney U test post hoc analysis (B). Genes were considered differentially expressed between groups if q-value ≤ 0.05. Categories of statistically significant differential gene expression are illustrated by bar graphs with the total number of genes written above for each category. Groups with different letters are significantly different. The majority of the differentially expressed genes were either upregulated (n = 808) or downregulated (n = 67) in WRF compared to C and W. **C**. Heatmap of standardized gene expression for genes upregulated or downregulated in WRF rats. The 18 animals used for each paradigm are sorted by the time of euthanasia. Higher levels of gene expression are displayed in yellow. **D**. Gene Ontology and KEGG pathways analysis of genes being significantly upregulated in the heart of WRF rats (n = 808). **E, F**. Left: Illustration of the 80S ribosome (E) and 26S proteasome (F), with upregulated genes in WRF rats *vs*. C and W rats being displayed in red. Genes whose expression is unaffected are displayed in grey. Right: expression profiles of ribosomal genes (E) and proteasomal genes (F) being upregulated in WRF rats. Each dot represents individual fpkm values. Groups with different letters are significantly different (p < 0.05, Kruskal–Wallis test).

Visualization of altered gene expression with heatmaps revealed biological variation between rats from the same group, indicating that rats were not equally affected by the different shift work protocols (Fig. 3C). However, individuals affected by shift work displayed a relatively homogenous transcriptional response to shift work, *i.e*., the impact of shift work on gene expression was observed for most genes (Fig. 3C). This suggests that shift work in WRF rats impacts entire transcriptional programs rather than random genes, and that some key master transcriptional regulators are affected by the lack of food intake during shift work.

To get insights into the transcriptional programs and biological pathways that were affected in the heart of WRF rats, we performed a KEGG pathway analysis using the 808 and 67 genes that were upregulated and downregulated in WRF rats, respectively. This analysis revealed a significant up-regulation of pathways associated with gene expression, including ribosome biogenesis, proteasome, spliceosome, and protein processing in the endoplasmic reticulum (Fig. 3D). In particular, 73 out of the 75 ribosomal protein genes that constitute the eukaryotic 80S ribosome showed increased expression in WRF rats (Fig. 3E). Similarly, 26 out of the 34 genes that make up the eukaryotic 26S proteasome displayed increased expression in WRF rats (Fig. 3F). Interestingly, genes upregulated in WRF were also enriched for pathways associated with the cell cycle, such as DNA replication, nucleotide excision repair, mismatch repair, and purine and pyrimidine metabolism. Together, these data suggest that changes in gene expression in the heart of WRF rats are associated with increased gene expression and cell division, a feature that may be related to cardiac hypertrophy as suggested in other studies (24-26).

The KEGG pathway analysis also revealed that genes exhibiting decreased expression in WRF rats were associated with various signaling pathways, including cholinergic synapse and adrenergic signaling in cardiomyocytes (Fig. 3D). While it is unclear why various signaling pathways are enriched for decreased gene expression in the heart of WRF rats, this may reflect the conflicting information between activity (*i.e*., work) and lack of food intake that WRF rats experienced.

### Altered gene expression in the heart of shift workers subjected to restricted feeding is associated with transcription factors known to regulate heart biology

The homogenous change in gene expression that was observed in each WRF rat (Fig. 3C) suggests that transcriptional programs driven by specific transcription factors are altered. Findings by the ENCODE consortium and others revealed that ∼95% of transcription factors bind in open chromatin regions, *i.e*., in genomic regions that are more sensitive to mild nuclease digestion (27-29). To get insights into the mechanisms underlying differential gene expression between WRF *vs*. C and W rats, we set out to uncover open chromatin regions in rat heart by carrying out DNase-Seq to identify the transcription factors that may be involved.

Analysis of our rat heart DNase-Seq dataset revealed that 28,514 genomic regions were more accessible to DNase I digestion, with a vast majority being located as expected in intergenic regions, transcription start sites, and introns (Fig. 4A, Table S5). To further assess the quality of our dataset, we performed a cross-comparison of heart DNase-Seq peaks between rat and mouse using mouse heart DNase-Seq datasets from the mouse ENCODE project (27). Using stringent peak calling, we found that that 71.2% of the rat DNase-Seq peaks overlapped with mouse DNase-Seq peaks (15,480 peaks out of 21,750 peaks) (Fig. 4B). Importantly, visualization of rat and mouse DNase-Seq profiles over syntenic genomic regions confirmed that many DNase hypersensitive sites (DHS, *i.e*., open chromatin regions) are conserved between rat and mouse, thereby validating our rat heart DNase-Seq dataset for further analysis of the transcription factors that contribute to the changes in gene expression in WRF rats (Fig. 4C).

**Fig. 4.**
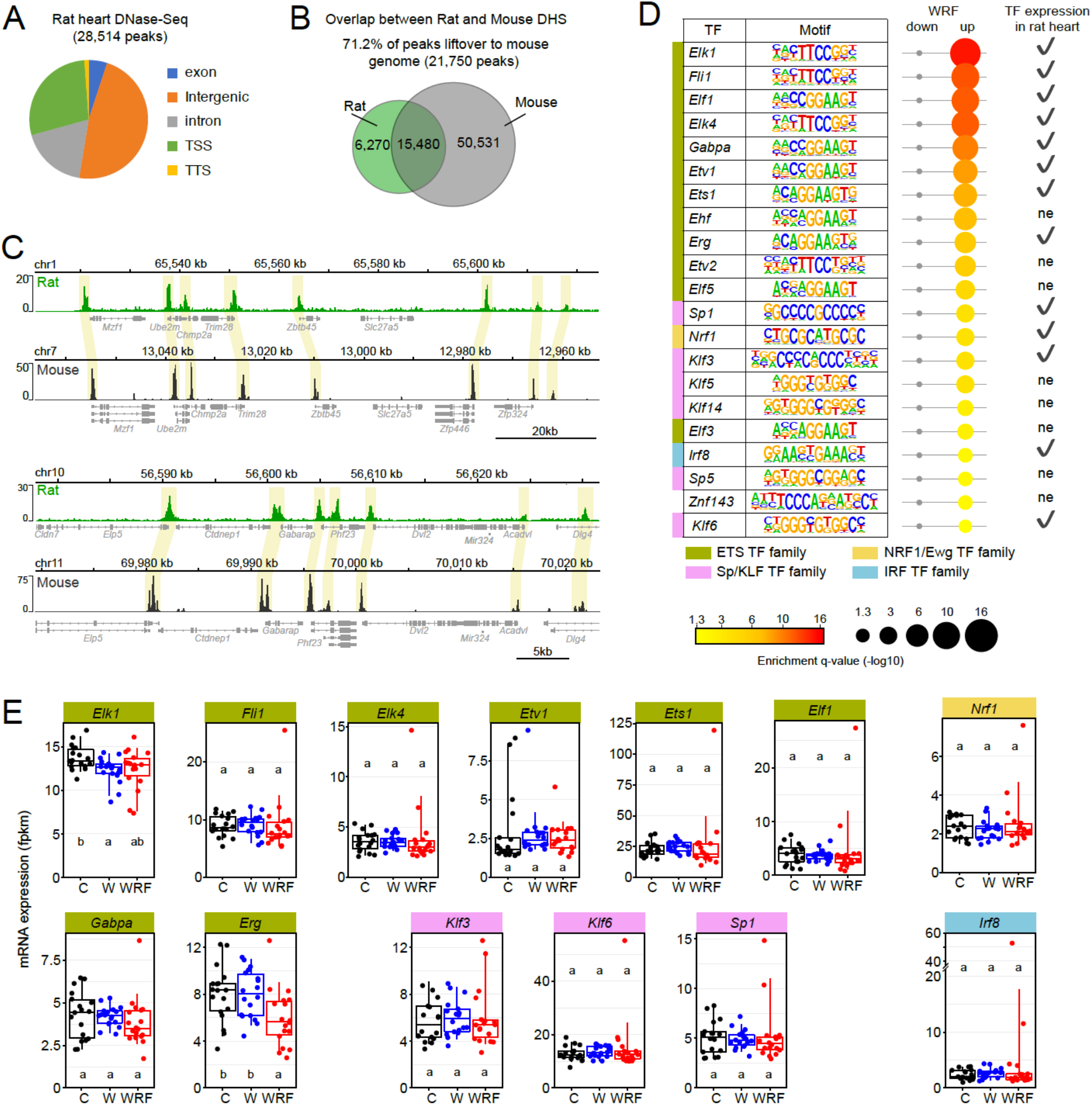
Transcription factor motif analysis at genes being misregulated in the heart of WRF rats. **A**. Genomic location of DNase I hypersensitive sites in the rat heart. **B**. Overlap between rat and mouse DNase-Seq peaks in the heart. Rat heart DNase-Seq peaks were converted to mouse mm10 genome with LiftOver and then overlapped with mouse heart DNase-Seq peaks (ENCODE project dataset). 76% of the lifted-over rat DHS overlapped with the mouse ENCODE dataset. **C**. IGV browser visualization of rat and mouse DNase-Seq reads at two syntenic genomic regions to illustrate the conservation of DNAse-Seq peaks between the two species. **D**. Motif analysis at DNase I hypersensitive sites located within genes being down- or up-regulated in WRF rats. Motifs with a q-value < 0.05 are colored from yellow to red with red being the most significant. Lack of enrichment for a motif (p > 0.05) is displayed in grey. Transcription factors expressed in the rat heart are illustrated with a checkmark and those that are not expressed in the rat heart are labeled ne. **E**. Expression profiles of several transcription factors whose motif is enriched in DNase-Seq peaks located in genes upregulated in WRF rats. Each dot represents individual fpkm values. Groups with different letters are significantly different (p < 0.05, Kruskal–Wallis test).

To identify these transcription factors, we conducted a motif analysis using HOMER in the open chromatin regions located within the 808 and 67 genes that were significantly increased and decreased in WRF rats, respectively (30). Several consensus motifs were identified, including those bound by the transcription factors belonging to the ETS family, the KLF family, NRF family, bHLH family, and for Ronin (Fig. 4D, Table S6). Interestingly, many of these transcription factors have been involved in the development of cardiovascular disorders and cardiac fibrosis (Table 1). For example, the ETS transcription factor *Ets1* has been shown to mediate angiotensin II-related cardiac fibrosis and cardiac hypertrophy (31, 32), while increased ETV1 activity has been shown to induce atrial remodeling and arrhythmia (33). Moreover, the SP/KLF family transcription factors *Klf3* and *Klf6* have been involved in cardiovascular development and associated with cardiac fibrosis, respectively (34, 35). In addition, the IRF family transcription factor IRF8 has been shown to suppress pathological cardiac remodeling by inhibiting calcineurin signaling (36) (Table 1).

**Table 1.**
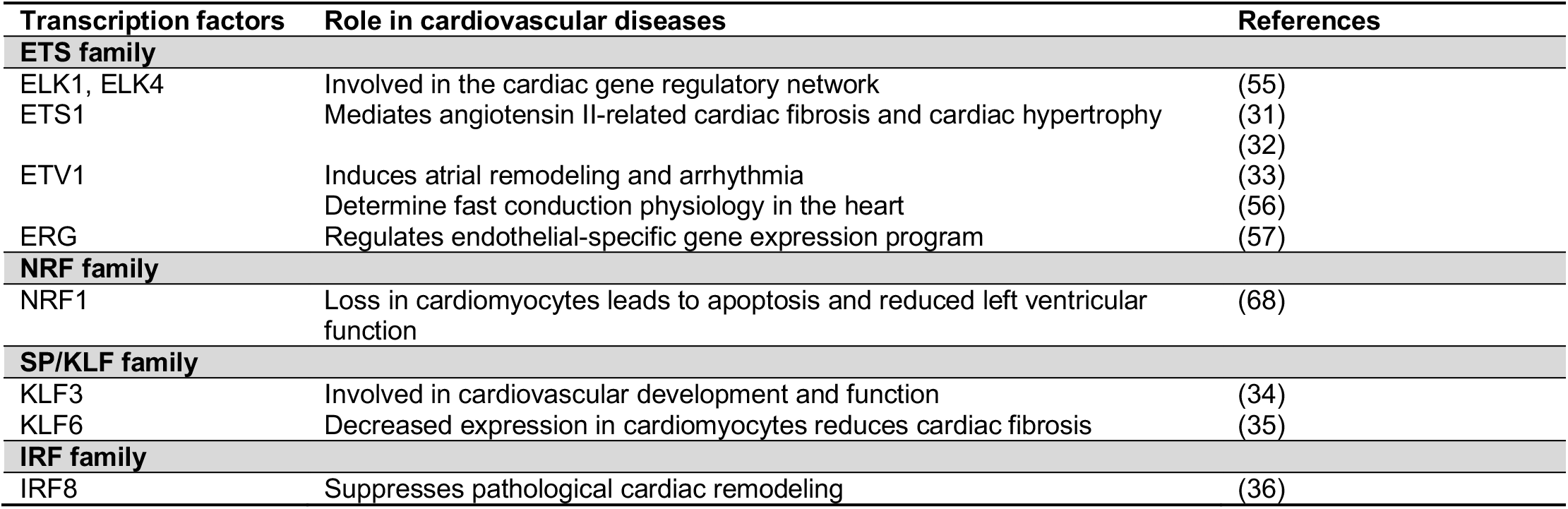
Transcription factors whose motif are enriched in genes up-regulated in WRF and their involvement in cardiovascular diseases.

Visualization of the transcript levels for these transcription factors in the heart of W, WRF, and C rats revealed that shift work protocols did not dramatically alter their expression (Fig. 4E). Only two of them, *Elk1* and *Erg*, showed decreased expression in W and WRF rats when compared to C rats (Fig. 4E). While we cannot exclude that the misregulation of gene expression in WRF rats is solely caused by *Erg* decreased expression in WRF rat heart, our data suggest that the lack of food intake during shift work alters heart gene expression by impairing the activity of several transcription factors (*e.g*., through mistimed activation of signaling pathways) rather than by affecting the expression of transcription factors.

### Genes whose expression is altered by shift work are associated with cardiac diseases

Several studies have characterized transcriptional signatures of cardiovascular disorders in both mice and humans. To determine whether genes being altered by shift work are also misregulated in pathological heart samples, we first examined a paper that identified cardiomyocyte gene programs being activated after cardiac hypertrophy and heart failure in the mouse (24). This study relied on transverse aorta constriction to promote pressure overload and induce cardiac hypertrophy 1-2 weeks after surgery and heart failure after 4-8 weeks. Using RNA-Seq of cardiomyocytes collected from 3 days to 8 weeks post-surgery, the authors clustered genes into 56 cardiac hypertrophy stage-specific modules that characterized distinct gene programs encoding morphological and functional transcriptional signatures occurring at specific stages of disease progression (Fig. 5A, S4) (24).

**Fig. 5.**
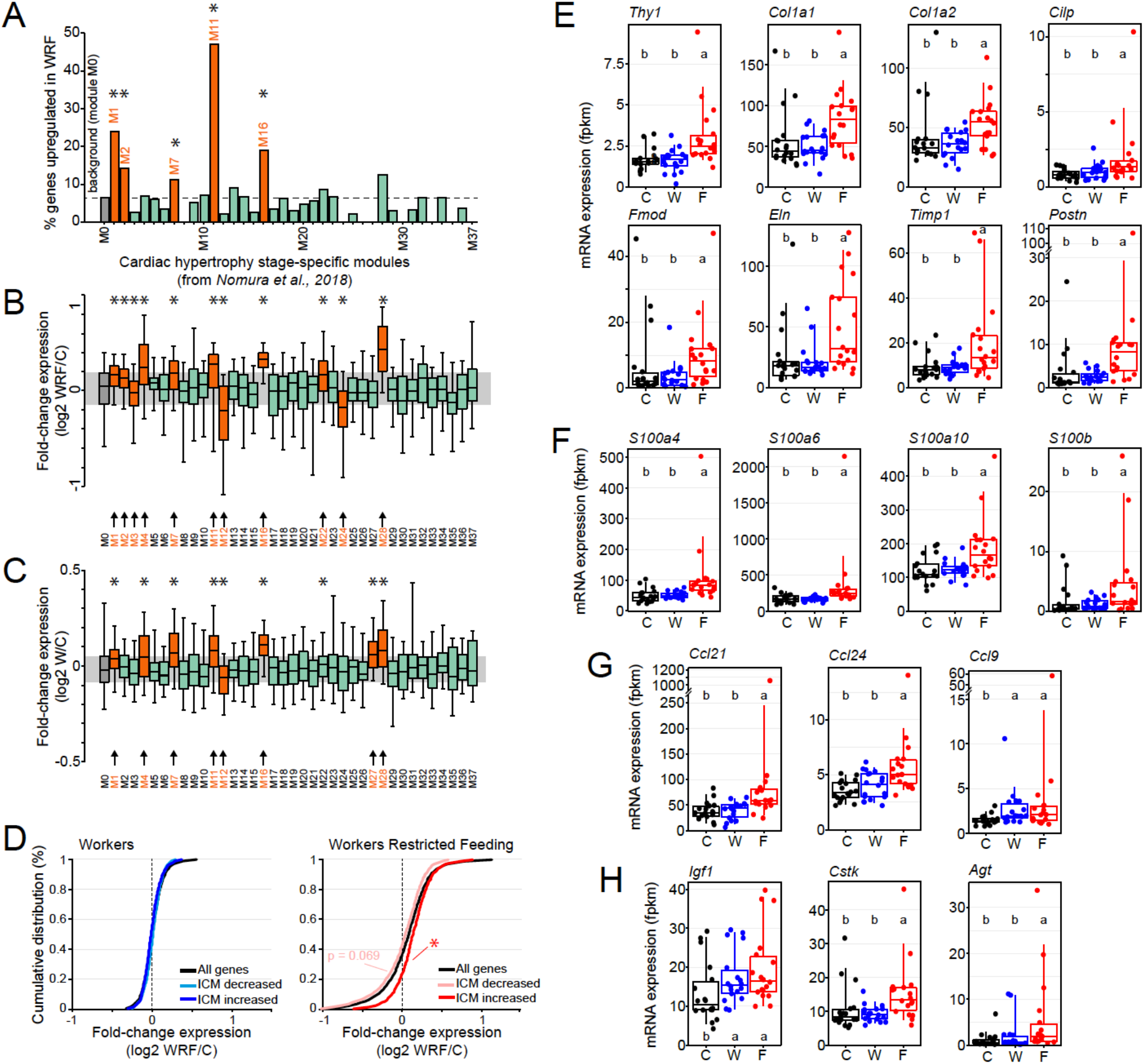
Association of genes misregulated in the heart of shift worker rats with heart diseases. **A**. Enrichment of genes upregulated in the heart of WRF rats for cardiac hypertrophy stage-specific modules. Modules were retrieved from Nomura et al., 2018, and correspond to distinct sets of genes being misregulated by cardiac hypertrophy in the mouse heart (24). Modules of at least 50 genes were considered in the analysis (M0 to M37). Module M0 comprises all genes unaffected by cardiac hypertrophy, while modules M1 to M37 correspond to functional sets of genes being affected at various stages of cardiac hypertrophy, *i.e*., from the early stage of cardiac hypertrophy to heart failure. Data are represented as the percentage of genes upregulated in the heart of WRF rats for each module. Modules showing a significant enrichment of genes are colored in orange and labeled with an asterisk (Fisher’s exact test; p < 0.05). **B**. Difference in gene expression between WRF and control rats for 38 modules described in Nomura et al., 2018 (24). The difference in expression was calculated for each gene as the WRF/control ratio (median of 18 samples for each group) and is represented in log2 scale. Modules colored in orange and labeled with an asterisk and arrow exhibit a significant difference in gene expression when compared to module M0, which comprises all genes unaffected by cardiac hypertrophy (Kruskal-Wallis test; p < 0.05). The grey area corresponds to the interquartile distribution of the control M0 module. **C**. Same analysis as B, but for W rats. **D**. Cumulative distribution of WRF/control or W/control expression ratio for all genes (black) or for genes described as either downregulated or upregulated by ischemic cardiomyopathy in the human heart. The list of down/up-regulated genes was retrieved from Sweet et al., 2018 (37). Distribution significant different from that of all genes is illustrated by an asterisk (Kolmogorov-Smirnov test, p < 0.05). **E-H**. Expression profiles of genes involved in the regulation of the extra-cellular matrix (E), of S100 family calcium-binding proteins (F), of chemokines (G), and other genes (H) whose misregulation has been implicated in heart diseases. Each dot represents individual fpkm values, and groups with different letters are significantly different (Kruskal-Wallis test; p < 0.05).

Restricting our analysis to modules of 50 genes or more (modules M0 to M37), we found that genes upregulated in the heart of WRF rats were enriched in five modules when compared to module M0, which comprised genes whose expression was unaffected by pressure overload (Fig. 5A, Table S7). Several of these modules (M1, M11, and M16) include genes encoding proteins involved in ribosomes, translation, oxidative phosphorylation, and generation of precursor metabolites and energy (Fig. S4, Tables S8, S9) (24). Interestingly, these modules were preferentially activated from 3 days to 2 weeks after transverse aorta constriction, *i.e*., early after pressure overload rather than at advanced hypertrophy and heart failure stages (24).

To extend our analysis, we examined changes in gene expression in WRF rat hearts for each module. While gene expression within most modules was unaffected and similar to the control M0 module, we found that genes within 11 modules were globally either upregulated (8 modules) or down-regulated (3 modules) (Fig. 5B, Table S10). Most of these modules were identical to those identified with the enrichment analysis, confirming that lack of food intake during shift work in rat reprograms cardiac gene expression to transcriptional signatures associated with pressure overload. Interestingly, the same analysis performed using changes in gene expression in shift worker rats (W group) revealed that gene expression within eight modules was also misregulated when compared to other modules (Fig. 5C, Table S11). Six out of eight modules (M1, M4, M7, M11, M16, and M27) were also upregulated in WRF rat heart, suggesting that shift work alone initiates changes in gene expression after 5 weeks, and that these effects are exacerbated if animals do not eat food during work.

We also examined transcriptional signatures associated with ischemic cardiomyopathy (ICM) and dilated cardiomyopathy (DCM) in humans, using public datasets comparing gene expression profiles between post-mortem pathological heart samples with non-failing heart samples (37). We found that genes upregulated in ICM hearts were globally upregulated in WRF rat heart, while those downregulated by ICM were slightly downregulated in WRF rat heart (Fig. 5D). The effects were less pronounced for genes being misregulated by DCM (Fig. S5), and no difference in gene expression profile was observed for shift worker rats (Fig. 5D). Taken together, these data indicate that lack of food intake during shift work induces changes in cardiac gene expression that resemble transcriptional signatures observed in pathological heart samples.

Closer inspection of the genes whose expression is altered in WRF rat heart revealed that many of them are associated with cardiovascular disorders. These include genes whose products are components of the extra-cellular matrix (ECM) such as collagen type 1 alpha 1 chain (*Col1a1)* and collagen type 1 alpha 2 chain (*Col1a2*), which are all upregulated in WRF rat heart (Fig. 5E). Because fibroblast markers including Thy-1 cell surface antigen (*Thy1*; Fig. 5E) and Fibroblast-specific protein-1 (*S100a4*; Fig. 5F) are also significantly upregulated in WRF rat heart, this suggests that lack of food intake during shift work may lead to collagen deposition and cardiac fibrosis. S100 proteins are calcium-binding proteins that regulate macrophage inflammation and whose expression is increased in the injured myocardium (38, 39). Given that three other members of the S100 family (Fig. 5F), and that several chemokines are also upregulated in WRF rats (Fig. 5G), this suggests that the heart of WRF rats is inflamed and colonized by immune cells. Finally, several genes whose increased expression has been shown to lead to cardiac hypertrophy and heart diseases are also upregulated in WRF rat heart, including *Insulin-like growth factor 1* (*Igf1*), *Angiotensin* (*Agt*), and *Cathepsin K* (*Ctsk*) (Fig. 5H) (40-43). Interestingly, the hormone *Igf1* and the inflammatory chemokine *Ccl9* are also upregulated in the heart of W rats, suggesting that shift work alone may also initiate some cardiac injury and inflammation in the heart (Fig. 5G, 5H).

### Shift work induces collagen deposition in the heart

Based on our findings that genes whose expression is altered by shift work are also misregulated in pathological heart samples from mice and humans, and since many of them regulate immune responses and ECM composition, we next examined if hearts from W and WRF rats displayed signs of inflammation and cardiac fibrosis.

First, we performed immunohistochemistry of two immune cells markers, CD45 and CD68, in the heart of W, WRF, and C rats. CD45 detects most cells of hematopoietic origin at all stages of maturation, while CD68 is a marker of inflammation associated with the involvement of monocytes/macrophages (44, 45). Consistent with the transcriptional response observed in WRF but not W rat hearts, the number of both CD45 and CD68 positive cells was significantly higher in the heart of WRF rats than that of W and control rats (Fig. 6A, 6B).

**Fig. 6.**
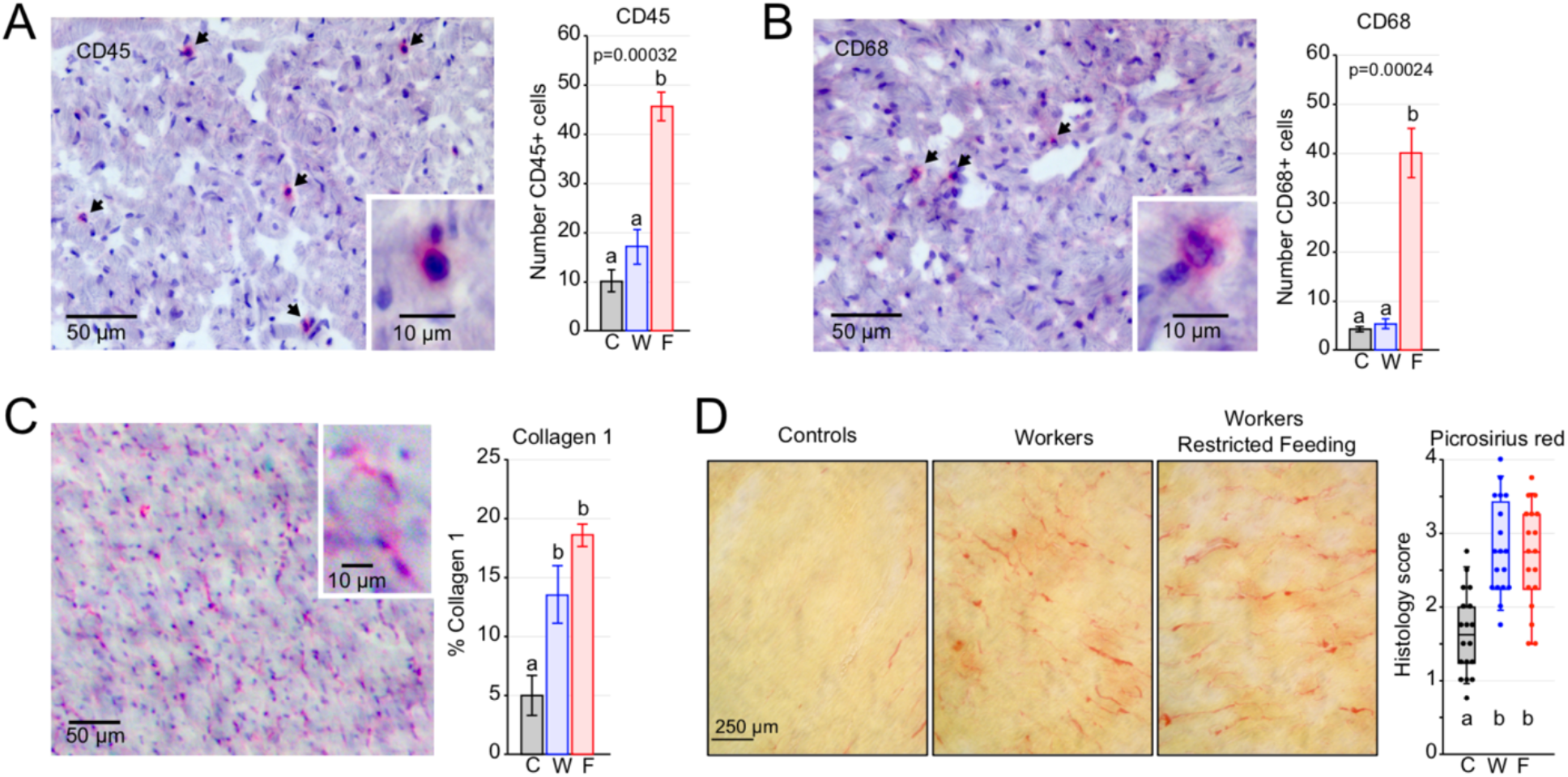
Inflammation and fibrosis in the heart of shift worker rats. **A, B**. Left: Illustration of CD45 (A) and CD68 (B) immunostaining in the heart of shift worker rats. Arrows indicate immuno-positive cells. Right: Quantification of the number of CD45+ (A) and CD68+ (B) in the heart of control, W, and WRF rats (n=3 per group). Groups with different letters are significantly different (one-way ANOVA; p < 0.05). **C**. Illustration of collagen 1 immunostaining (left) and quantification of collagen 1 signal (right) in the heart of control, W, and WRF rats (n=3 per group). Groups with different letters are significantly different (one-way ANOVA; p < 0.05). **D**. Left: representative picrosirius red staining in the heart of control, W, and WRF rats. Right: quantification of the picrosirius red staining (n=18 per group). Quantification was performed using a scale from 0 to 4 (0 = no staining, 4 = maximum staining; see Fig. S5). Each dot corresponds to the average score given by four investigators for each sample. Groups with different letters are significantly different (one-way ANOVA; p < 0.05).

Next, we performed immunohistochemistry of Collagen 1 to determine whether shift work leads to cardiac fibrosis. Surprisingly, the hearts of both W and WRF rats showed increased Collagen 1 expression (Fig. 6C), *i.e*., W rat heart exhibited signs of fibrosis despite any increase in *Col1a1* and *Col1a2* expression (Fig. 5E). To confirm this result, we conducted picrosirius red staining of all C, W, WRF rat hearts (n = 18 per group), and double-blind scored collagen deposition based on a 0 to 4 scale (0: no collagen deposition; 4: maximum collagen deposition; Fig. S6). Analysis of picrosirius red staining confirmed Collagen 1 immunohistochemistry, *i.e*., collagen deposition was significantly higher in W and WRF hearts compared to C rat hearts (Fig. 6D).

Taken together, these results indicate that while lack of food intake during shift work (but not shift work alone) leads to increased cardiac inflammation, five weeks of shift work alone is sufficient to promote cardiac fibrosis. They also suggest that shift work induces some heart damages and that these effects are potentiated if animals do not have access to food during work.

## Discussion

Several epidemiological studies have shown that shift workers have an increased risk of developing cardiovascular diseases (11, 13-15). However, the underlying mechanisms are for the most part unknown as no study has directly investigated in an animal model how shift work affects the heart biology and physiology. Using a well-characterized model of shift work in rats (22), we provide evidence that shift work reprograms the heart cycling transcriptome and leads to cardiac fibrosis. Importantly, our data also demonstrate that lack of food intake during shift work, an intervention that prevents obesity and diabetes in rat shift workers, does not preclude cardiac fibrosis. Restricting food access during shift work indeed markedly disrupts gene expression in the heart, and leads to cardiac fibrosis and inflammation. Together, our findings provide insights on how shift work affects cardiac functions, and strongly suggest that strategies aiming at mitigating the development of metabolic disorders in shift workers may have adverse effects on cardiovascular diseases.

Our genome-wide analysis of gene expression revealed that shift work reprograms the heart cycling transcriptome. In shift worker rats, the number of rhythmic genes is decreased by half and the molecular clock oscillation is impaired. Intriguingly, the effect on clock gene rhythmic expression is primarily due to differences between replicated shift work protocols rather than a global dampening of clock oscillations in individual rats (Figure S1B). As shown previously, shift work in rats shifts the hepatic clock rhythms due to the reversal of the food intake rhythm, *i.e*., because shift workers eat preferentially during their work shift (20, 22). Given that peripheral clocks can be synchronized by multiple cues including rhythms in food intake, hormones, body temperature, and neuronal activity, the heart clock may be less sensitive to changes in rhythmic food intake than the liver clock, and more sensitive to other cues such as rhythms in locomotor activity or in neuronal activity that originate from the SCN (22). This possibility is supported by findings showing that PER1 rhythmic expression is shifted by shift work in some hypothalamic structures (*e.g*., arcuate nucleus, dorsomedial hypothalamus), but not in others (*e.g*., SCN, paraventricular nucleus) (46). However, why replicate rhythms did not respond homogenously still remain unclear, and warrant further investigation in future experiments.

The diversity in cues that can synchronize peripheral clocks may also explain why most clock genes exhibit dampened rhythmic expression in WRF rat hearts (Fig. 2D, S1A). Indeed, the synchronizing effect of rhythmic food intake may be counteracted by antiphasic cues driven by the sleep/wake cycle, thereby reducing the amplitude of the circadian clock oscillation (47).

Interestingly, although the molecular clock oscillations are dampened in WRF rat hearts, the number of rhythmic genes is two-fold higher than in control rat hearts. Since rhythmic gene expression can be driven by feeding rhythms independently of the molecular clock (48-50), this increased rhythmic gene expression in the heart of WRF rats is likely caused by the 8 hours of fasting occurring during work. This would explain why rhythms in gene expression occur at discrete phases rather than across the 24-hour day. Consistent with this possibility, the expression of many genes involved in oxidative phosphorylation is decreased at the end of shift work, likely reflecting a decrease in substrates for the generation of ATP. Moreover, genes involved in endocytosis are increased during shift work, maybe as a result of compensatory mechanisms aimed at increasing the uptake of extracellular molecules for energy sources (51).

In addition to dampening the molecular clock oscillations and reprogramming the cycling heart transcriptome, lack of food intake during shift work strongly impairs gene expression in the heart. Because these effects are less pronounced in shift worker rats, impaired gene expression in WRF rat hearts is likely due to the conflicting information between the work schedule, where animals are awake and likely to have increased heart workload due to increased sympathetic neural activity (52), and the lack of food intake. Motif analysis in DNase-Seq peaks located in misregulated genes in WRF rats revealed that transcription factors of the ETS family are likely involved (Fig. 4D). ETS transcription factors are key regulators of cell proliferation, differentiation, angiogenesis, and inflammation, and their transcriptional activity is regulated by signaling pathways including the MAP kinase pathway (53, 54). Interestingly, our KEGG analysis pathway revealed that components of the MAPK signaling pathway are rhythmically expressed in C rats but not in WRF rats, and that they are also downregulated in the heart of WRF rats (Fig. 2E, 3D). Importantly, ETS transcription factors contributes to many aspects of the heart physiology, and alteration of their transcriptional activity is associated with cardiovascular diseases including cardiac fibrosis and arrhythmia (31-33, 55-57). Other transcription factor motifs were identified, including SP/KLF family members and IRF8. Interestingly, these transcription factors have also been involved in the regulation of heart physiology and for some of them, shown to promote cardiac diseases when their activity is impaired (Table 1).

Several biological pathways that are upregulated in WRF rat hearts are involved in protein synthesis and degradation, DNA synthesis, and gene expression (*e.g*., purine and pyrimidine metabolism). Interestingly, these processes have been shown to be increased in the early stages of cardiac hypertrophy, suggesting that five weeks of shift work without food intake during work impairs heart functions (24, 26, 58). Consistent with this possibility, many genes misregulated in WRF rat heart are also misregulated in cardiomyocytes several days after transverse aorta constriction, *i.e*., when the heart is under pressure overload and in the early stage of cardiac hypertrophy, and before its progression to heart failure (24). Moreover, many genes upregulated in WRF rat hearts are known to be misregulated in ailing hearts (Fig. 5E-H). For example, increased expression of *Col1a1* and *Col1a2*, which encode for components of type I collagen, has been linked to remodeling of the cardiac matrix (59), and upregulation of collagen 1 is a well-established marker of cardiac fibrosis and is observed after cardiac hypertrophy (60-62). This upregulation of *Col1a1* and *Col1a2* expression, along with other markers of cardiac fibrosis like *Thy1, Postn*, and *Cilp* (63-65), is consistent with our findings that the heart of WRF rat is fibrotic (Fig. 6C, 6D). In addition, the increased expression of several chemokines and members of the S100 protein family, along with the increased number of CD45+ and CD68+ cells, also indicate that the heart of WRF rats is inflamed and colonized by immune cells. Given that inflammation of the myocardium also occurs after hypertrophic cardiopathy (66, 67), our data altogether strongly suggest that lack of food intake during shift work is associated with cardiac hypertrophy.

While our initial RNA-Seq analysis did not identify large defects in gene expression in the heart of W rats, our analysis of modules associated with pressure overload, cardiac hypertrophy, and its progression to heart failure uncovered significant differences between shift worker and control rats (24). This discrepancy is likely due to our stringent analysis of differential gene expression, which comprised a Benjamini-Hochberg p-value correction because of the multiple comparisons (Fig. S3). Most modules exhibiting impaired gene expression in W rats were also found in WRF rats, suggesting that shift worker rats were in the early stage of developing cardiac diseases. Consistent with this possibility, hearts of shift worker rats displayed cardiac fibrosis, but no sign of inflammation. While it still remains unclear why W rat hearts exhibit increased collagen deposition without changes in *Col1a1* and *Col1a2* mRNA expression, their expression may be acutely upregulated by shift work, i.e., their expression is similar to those observed in C rats during weekends when the tissues were collected. Future experiments will be required to test this possibility.

In summary, our results indicate that five weeks of shift work in rats are sufficient to promote cardiac fibrosis, whereby the effects of shift work are exacerbated when animals do not have access to food during work. Based on these data, it is tempting to speculate that the differences we observed between W and WRF shift worker rats may be relevant to explain some of the heterogeneity that has been reported in epidemiological studies of human shift workers (10, 12). Importantly, our findings also suggest that lack of food intake during work, which was proposed as a possible intervention to prevent the development of metabolic disorders in shift workers, may not be appropriate to prevent the development of cardiovascular diseases in shift workers.

## Methods

### Animals

Seven-to-eight-week-old male Wistar rats were exposed to a shift work protocol for five weeks. Each animal was housed individually in a 12 L:12 D cycle (7 am to 7 pm) throughout the protocol (Figure 1). The protocol consisted of 54 animals randomly assigned to three groups: 18 Controls ‘C’ (undisturbed and no shift work protocol, *ad libitum* food and water), 18 Workers ‘W’ (shift work protocol, *ad libitum* food and water), 18 Worker Restricted Feeding ‘WRF’ (shift work protocol, *ad libitum* water, access to food only outside shift work). Shift work protocol consisted of five days of scheduled ‘work’ (workday: Monday to Friday, 9 am to 5 pm) followed by two days without ‘work’ and given *ad libitum* food and water (weekend: Saturday and Sunday). Experiments were approved by Universidad Nacional Autonoma de Mexico, in accordance with animal handling, Norma Oficial Mexicana NOM-062-ZOO-1999.

### Shift work protocol

To simulate shiftwork, rats were placed in a slowly rotating drum that was designed to keep the animal awake during their natural rest phase, as previously described (22). Briefly, drums rotated with a speed of one revolution per three minutes, and enabled animals to move, eat, and drink freely. However, animals were not able to fall asleep and had to stay awake without requiring them to exhibit laborious movements. Food and water were hung from a concentric middle tube and freely available, unless noted otherwise for the WRF group. Outside of work hours, rats were housed in their individual cages with food and water provided *ad libitum*, and kept under 12:12 LD cycle.

### Heart Tissue Collection

After five weeks of shift work protocol, rats were euthanized on a non-working day schedule every four hours starting one hour after light on (ZT01) at 7 am. Hearts were collected, flash-frozen in liquid nitrogen, and shipped to Texas A&M University.

### Whole transcriptome sequencing (RNA-Seq)

RNA was isolated from crushed frozen heart samples using Trizol reagent according to the manufacturer’s instructions (Life Technologies). Total RNA was diluted to 4 μg, and subjected to polyA selection using Dyna oligo dT beads following manufacturer’s instructions (Invitrogen). Sequencing libraries were constructed using NEB Ultra Directional RNA Library Prep Kit from Illumina (New England BioLabs) following manufacturer’s instructions. Briefly, polyA purified RNA was fragmented for 15 minutes at 94°C before first and second-strand cDNA synthesis. Purification of cDNA was done using AMPure XP beads following NEB Ultra Directional RNA Library Prep Kit instructions. Libraries were then PCR-amplified for 13 cycles, bead-purified, and quantified using a Quantus Fluorometer (Promega) and qPCR. Libraries were generated with multiple bar-coded adaptors, mixed in equimolar ratio, and sequenced on three lanes of an Illumina NextSeq 500 with a sequencing length of 76bps.

### Rat heart DNase-Seq

Nuclei were isolated from crushed frozen heart samples of control C rats (n = 2 biological replicates) using a dounce homogenizer in lysis buffer (0.32 M sucrose, 10 mM Tris-HCl pH 8, 5 mM CaCl2, 5 mM MgCl2, 2 mM EDTA, 0.5 mM EGTA, 1 mM DTT). Samples were homogenized 6-times with pestle A and 4-times with pestle B. Homogenates were mixed with sucrose lysis buffer (2.2 M sucrose, 10 mM tris HCl pH 8, 5 mM CaCl2, 5 mM MgCl2, 2 mM EDTA, 0.5 mM EGTA, 1 mM DTT) and then layered on top of a sucrose cushion (2.05 M sucrose, 10% glycerol, 10 mM HEPES pH7.6, 15 mM KCl, 2 mM EDTA, 1mM PMSF, 0.15 mM spermine, 0.5 mM spermidine, 0.5 mM DTT). Samples were ultracentrifuged for 45 min at 24,000 rpm (100,000 g) at 2°C in a Beckmann SW28 rotor. The supernatant was removed and nuclei were washed three-times in resuspension buffer (10 mM Tris, pH 7.5, 150 mM NaCl, 2 mM EDTA, 1 mM PMSF). Nuclei were then counted, aliquoted (five million nuclei per tube) and flash-frozen. On the day of DNase digestion, nuclei were thawed, pelleted, and digested for 3 minutes at 37°C in DNase I digestion buffer (6 mM CaCl2, 78.5 mM NaCl, 13.5 mM Tris-HCl pH 8, 54 mM KCl, 0.9 mM EDTA, 0.45 mM EGTA, 0.45 spermidine) supplemented with 4 µl of DNase I (80 units/mL). After digestion, an equal volume of stop buffer (50 mM Tris-HCl pH 8, 100 mM NaCl, 0.1% SDS, 100 mM EDTA, 1 mM spermidine, 0.3 mM spermine) was added to each tube and incubated at 55°C for 1 hour. RNA was digested by adding 4.5 µl of RNase A (10 mg/mL) and samples were incubated at 37°C for 30 minutes. Samples were purified by phenol/chloroform extraction and resuspended in water. DNA size fractionation was done using a sucrose gradient and centrifuged for 30,000 rpm for 20 hours at 20°C in an SW-41 rotor with no brake and minimum acceleration. DNA fractions with fragment size under 500 bp were pooled together, ethanol purified, and resuspended in water. DNase-Seq libraries were generated using NEBNext® ChIP-Seq Library Prep Master Mix Set NEB) as per the manufacturer’s instructions. DNA collected from the DNase I digestion was quantified using a Quantus Fluorometer (Promega), and 10 ng was used to generate the libraries. Libraries were PCR-amplified for 14 cycles using Phusion Taq (M0530S). Adaptor dimers were removed using gel purification, and libraries were purified by phenol-chloroform extraction followed by ethanol precipitation. Libraries were quantified using a Quantus fluorometer and qPCR, and subjected to paired-end sequencing (2 × 75bp) using an Illumina NextSeq 500.

### Alignment of RNA-Seq libraries and differential gene expression analysis

Libraries were sequenced to a median depth of ∼25 million reads/sample and mapped to the *Rattus norvegicus* Rn6 genome assembly using Tophat2 alignment and the following criteria: -- read-realign-edit-dist 2 -g 1 --b2-sensitive (69). Sequencing depth and percentage alignment are provided in Table S12. Gene expression was analyzed using Cufflinks with the following criteria: --library-type fr-firststrand (70). Visualization of the forward and reverse strands were done using a custom shell script with total signal normalized to 10 million reads and viewed with the Integrated Genome Viewer (71). Only genes expressed above a fpkm threshold of 1 in at least 18 samples out of the 54 total samples were considered for statistical analysis (n = 10,257 genes). Differential gene expression was performed using a Kruskal-Wallis test with Benjamini-Hochberg correction for multiple comparisons followed by a Mann-Withney U test post hoc analysis to identify differences between groups. Genes were considered to be expressed differently between group if q ≤ 0.05.

### Alignment of DNase-Seq libraries and motif analysis

Libraries were sequenced to a median depth of ∼ 36 million and mapped to the *Rattus norvegicus* Rn6 genome with bowtie2 using the following criteria: -t –phred33 -X 650 (72). The average alignment was of ∼ 85%. Sam files were converted to bam files and merged using SAMtools view, sort, and merge (73). Duplicates were removed with Picard 2.8.2 MarkDuplicatesWithMateCigar created by the Broad Institute. A custom python script was used to remove unpaired reads, unmapped reads, and filled in sequences between paired-end reads. Sorted coordinates were then made into BWfiles using Bedtools, normalized to 10 million reads, and viewed with the Integrated Genome Viewer (71, 74). Peak calling was done using findPeaks from the HOMER suite using the following commands: -size 45 -ntagThreshold 2 - region (30). Open chromatin regions for genes of interest were determined by using HOMER gene annotation and annotation for these genes were extended 10kb from transcription start sites and 1kb from transcription termination sites. These gene windows were overlapped with DNase-Seq reads using intersectBed from Bedtools suite (74). Motif analysis was done on these overlapping regions using the perl script findMotifsGenome.pl from HOMER using the following command: -size given (30).

### Analysis of rhythmic gene expression

Rhythmicity analysis was performed using six algorithms from four programs: F24 (75, 76), JTK_CYCLE, ARSER, and LS from MetaCycle (77), HarmonicRegression (78), and RAIN (79). The resulting p-values from all 6 algorithms were combined using Fisher’s method into one p-value, which was then adjusted to control for the false-discovery rate (FDR) using the Benjamini-Hochberg method (80) within the p.adjust function available in base R. Genes with a q-value under 0.05 were considered as rhythmically expressed.

### KEGG pathway and Gene Ontology analyses

All KEGG and GO analyses were performed using the kegga and goana functions available within the R package Limma (81). Gene symbols were converted to Entrez IDs with AnnotationDbi (82).

### Effects of shift work on cardiac hypertrophy stage-specific modules

The identity of the genes clustered into the 56 cardiac hypertrophy stage-specific modules was retrieved from Nomura et al., 2018 (Supplementary Data 4; 24), and cross-referenced with our list of 10,257 genes expressed in the rat heart. To avoid statistical bias due to low number of genes in a module, we restricted our analysis to modules of 50 genes or more (modules M0 to M37). Enrichment was calculated as the percentage of genes being upregulated in the heart of WRF rats for each module, and considered significant if p ≤ 0.05 when analyzed with a Fisher’s exact test. Differences in gene expression between WRF and C rats (Figure 5B) and between W and C rats (Figure 5C) were calculated for each gene as the WRF/control ratio (median of 18 samples for each group). Differences in gene expression between the control module M0 and the modules M1 to M37 were analyzed using a Kruskal-Wallis test and considered significant if p ≤ 0.05.

### Effects of shift work on the expression of genes being misregulated by ischemic cardiomyopathy and dilated cardiomyopathy

The list of genes being down/up-regulated in the heart of patients suffering from ischemic cardiomyopathy (ICM) and dilated cardiomyopathy (DCM) was retrieved from Sweet et al., 2018 (Table S3; 37). This list was then cross-referenced with our list of 10,257 genes expressed in the rat heart. Differences in the distribution between all genes and genes either up/down-regulated by ICN and DCM were assessed using Kolmogorov-Smirnov test, and considered significant if p < 0.05. Distributions were normalized to 500 quantiles to reduce the effect of size on the Kolmogorov-Smirnov test sensitivity (83).

### Immunohistochemistry

Immunohistochemistry was performed as previously described with some minor modifications (84, 85). Serial 10 µm thick cryosections of the heart were air-dried overnight and fixated in fresh acetone for 15 minutes at room temperature. Slides were then air-dried for 10 minutes and hydrated for 5 minutes in water and 5 minutes in PBS. Nonspecific binding was then blocked by incubation with PBS containing 2% BSA (fraction V, globulin free; Sigma-Aldrich) for 30 minutes. Endogenous biotin was blocked by the addition of streptavidin and biotin solutions, following the manufactures’ instructions (Streptavidin/Biotin Blocking Kit, Vector Laboratories, Burlingame, CA). Slides were then incubated with primary antibodies, anti-CD45 (5 µg/mL, MCA340R, OX-33, mouse IgG1, Bio-Rad), anti-CD68 (5 µg/mL, MCA341R, ED1, mouse IgG1, Bio-Rad), or anti-Collagen 1α1 (1 µg/mL, NB600-408, rabbit IgG, Novus Biologicals) in 0.5% NP40 and 0.05% SDS overnight at 4°C. Isotype-matched irrelevant mouse monoclonal antibodies or irrelevant polyclonal antibodies (BD Biosciences, or Jackson ImmunoResearch Laboratories) were used as controls. Slides were then quickly rinsed twice in PBS, and then 3 × 5 min in PBS. Primary antibodies were then detected with biotin-labeled F(ab’)2 anti-mouse IgG (Jackson ImmunoResearch Laboratories) or anti-rabbit IgG (Southern Biotechnology Associates) in PBS containing 2% BSA, as appropriate for 30 minutes. Slides were quickly rinsed twice in PBS, and then 3×5 min in PBS, before being incubated for 30 minutes with 1/500 ExtraAvidin-Alkaline Phosphatase (Sigma-Aldrich) in PBS containing 2% BSA. Slides were then rinsed with PBS and incubated for 10 minutes with 200 mM TRIS pH 8.3. Staining was then developed with Vector Red Alkaline Phosphatase Kit (Vector Laboratories) for 10 min. Sections were then counterstained for 1 min with Gill’s hematoxylin No. 3 diluted 1/5 with water (Sigman-Aldrich), and were then rinsed in water. Slides were dehydrated through 95% and 100% ethanol; cleared with xylene, and mounted with Permount (VWR).

The number of positive cells was assessed by two individuals counting 10 blinded selected fields per section with a random start position. Collagen was analyzed using ImageJ software using the standard algorithms to define the areas of staining (Rasband, W. S., ImageJ U.S. National Institute of Health, Bethesda, MD).

### Picrosirius red staining

Characterization of fibrosis was performed using picrosirius red, which specifically binds to collagen, on all 54 heart samples as previously described (84, 85). Heart samples were embedded in Tissue-Tek OCT, sliced 10 µm thick with a cryostat and slices were air-dried to slides. Slides were stained in picrosirius red for one hour, washed in acidified water twice, and dehydrated in three changes of 100% ethanol. Slides were then cleared with xylene and mounted with a resinous medium.

Quantification was performed using a scale from 0 to 4 (0 = no staining, 4 = maximum staining) as shown in Figure S5, by four investigators double-blinded to the conditions. The final score for each animal corresponds to the average score given by all four investigators.

### Data availability

Sequencing datasets have been deposited on Gene Expression Omnibus (GSE124870) and can be accessed using the following link: https://www.ncbi.nlm.nih.gov/geo/query/acc.cgi?acc=GSE124870 Access to the datasets is currently restricted, but can be requested to the corresponding author.

## Supporting information

Trott_Tables_S1-S8,S10-S12

Trott_Table_S9

## Acknowledgements

We are grateful to Cynthia Córdoba-Manilla for her assistance with the shift work protocol and tissue collection. We are thankful to Michael Rosbash, Kate Abruzzi, and Ryanne Spann for helping with the sequencing of the RNA-Seq libraries. We also thank the Texas A&M Institute for Genome Sciences and Society for helping with sequencing the DNase-Seq libraries and for the maintenance of their high-performance computing cluster. We are grateful to Matt Sachs for helping with setting up the DNase-Seq technique, and to Darrell Pilling for helping with immunohistochemistry. This work was supported by startup funds from Texas A&M University, a Texas A&M University-CONACYT Research Grant (2015-033), and in part by the National Institutes of Health (R21 AI144454).

## Authors contributions

A.J.T., R.M.B., and J.S.M. conceived and designed the experiments. N.N.G.V., C.E., and R.M.B. performed the shift work experiment and collected the heart samples. A.J.T. generated the RNA-Seq and DNase-Seq libraries, and performed the picrosirius red staining. A.J.T. and T.R.K. performed the immunohistochemical assays. A.J.T., B.J.G., and J.S.M. performed the bioinformatics analysis. A.J.T. and J.S.M. wrote the manuscript, and all authors contributed to editing the final draft.

## Supplementary Tables

Table S1: Fpkm data, analysis of rhythmicity and gene expression levels (related to Figures 2, 3, and S3).

Table S2: KEGG pathway analysis, rhythmically expressed genes (related to Figure 2)

Table S3: List of genes being rhythmically expressed in each enriched KEGG pathway (related to Figure 2 and S2)

Table S4: KEGG pathway analysis, differentially expressed genes (related to Figure 3)

Table S5: List of DNase I hypersensitive sites in rat heart and genomic location (related to Figure 4)

Table S6: Motif analysis for genes being up-regulated or down-regulated in WRF rats (related to Figure 4)

Table S7: Results statistical analysis for figure 5A (related to Figure 5)

Table S8: KEGG pathway analysis of the genes assigned to the modules M0 to M37 (related to Figure 5)

Table S9: Gene Ontology analysis of the genes assigned to the modules M0 to M37 (related to Figure 5)

Table S10: Results statistical analysis for figure 5B (related to Figure 5)

Table S11: Results statistical analysis for figure 5B (related to Figure 5)

Table S12: Summary of RNA-Seq library size and percentage alignment (related to Methods)

**Fig. S1.**
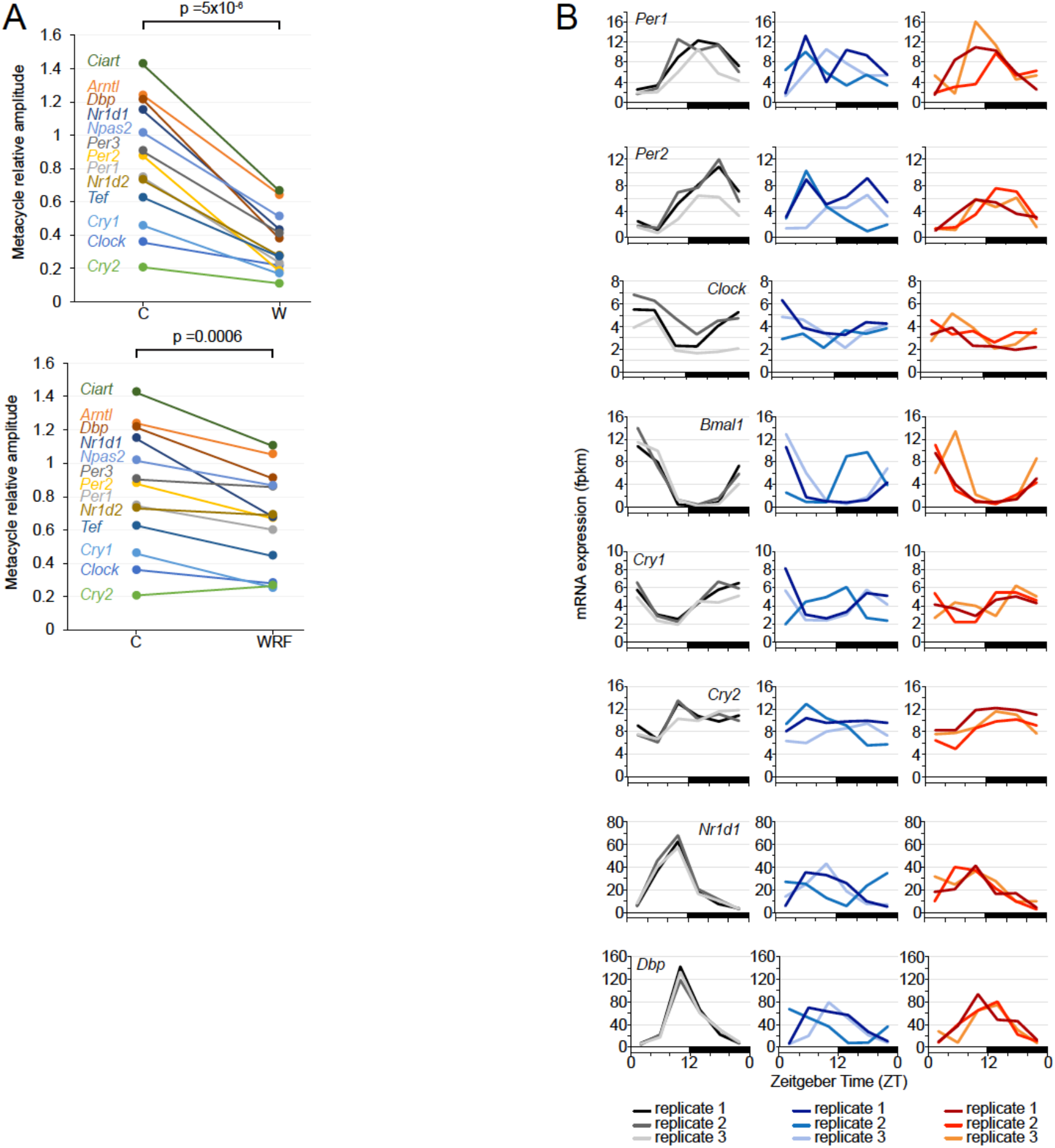
13 core clock genes in the heart of C, W, and WRF rats. The relative amplitude was calculated using Metacycle, and differences between groups was assayed using a two-tailed paired student t-test. **B**. Expression of eight core clock genes in the heart of C, W, and WRF rats. The expression is displayed for each of the three independent replicate rhythms for each group. Clock gene peak expression in replicate 2 of W rats is in antiphase to all other replicates.

**Fig. S2:**
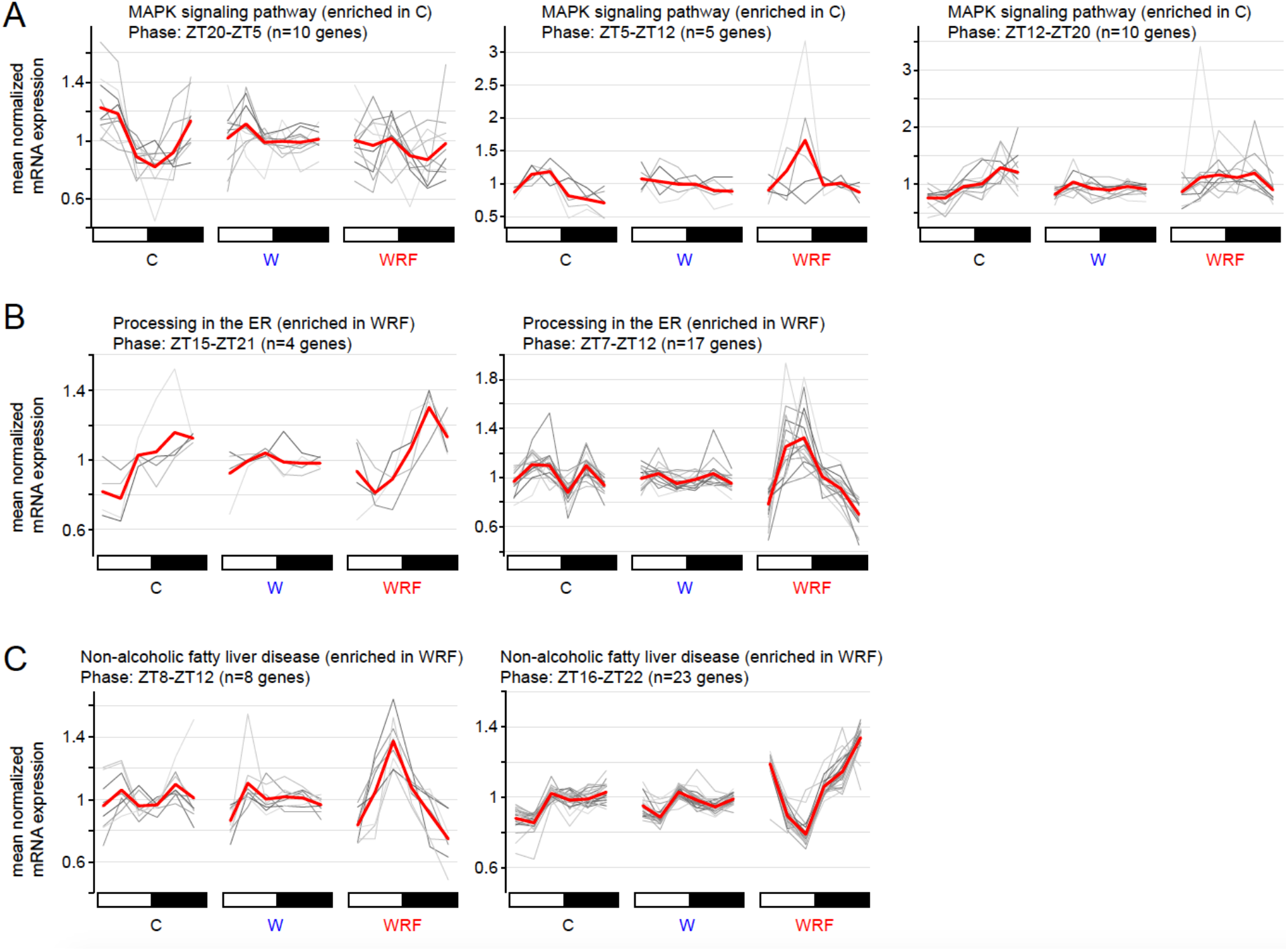
Effect of shift work on the rhythmic expression of genes accounting for enriched KEGG pathways. Expression profile of genes accounting for the enrichment of three KEGG pathways: MAPK signaling pathway, enriched in C rats; Processing in the ER, enriched in WRF rats; and Non-alcoholic fatty liver disease, enriched in WRF rats. The expression of individual genes is displayed in grey, and the averaged expression in red. For each enriched pathway, gene expression is parsed based on the phase of rhythmic gene expression.

**Fig. S3:**
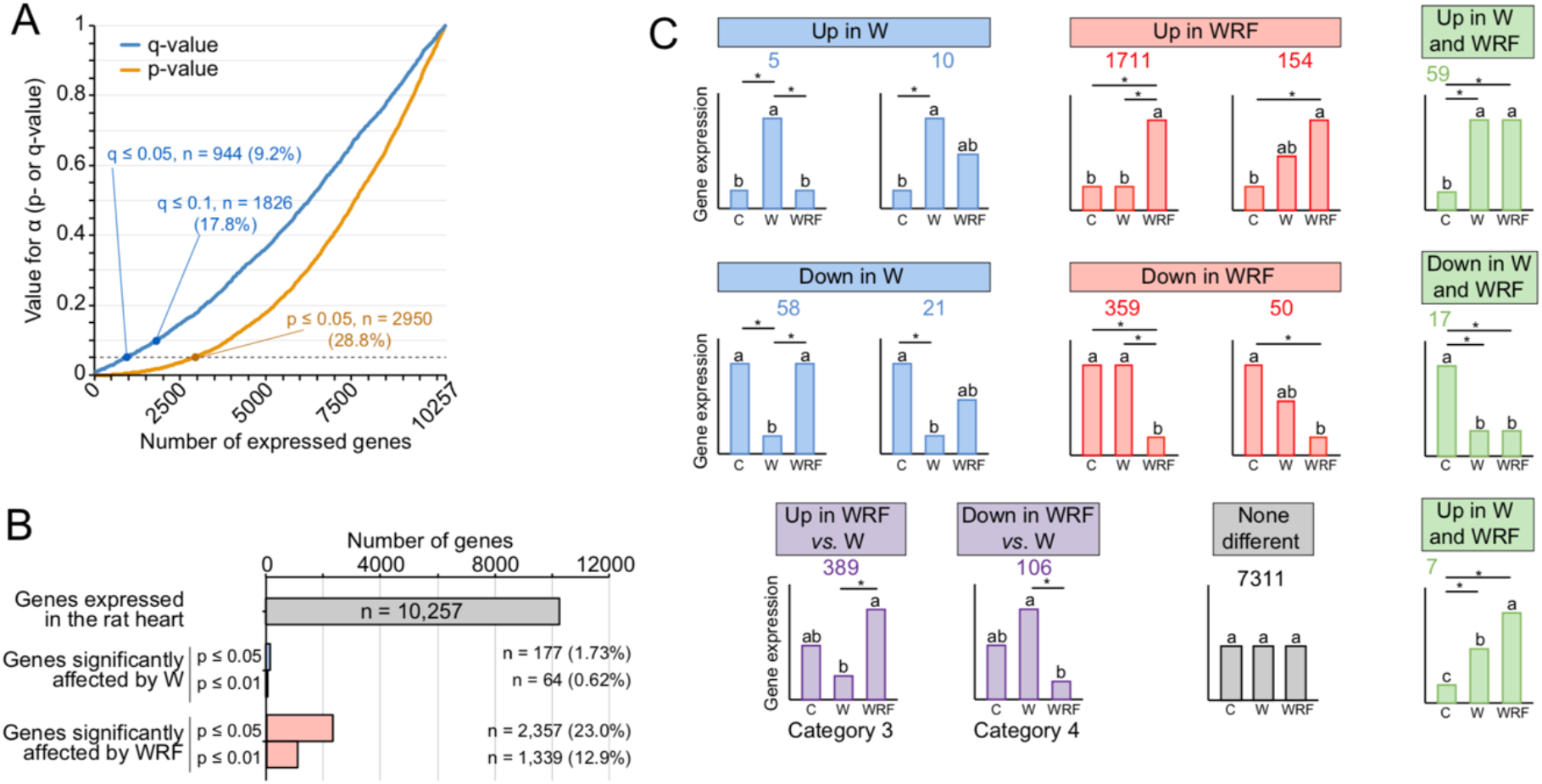
Analysis of differential gene expression between C, W, and WRF rats. Analysis of differential gene expression between C, W, and WRF rat hearts was conducted using a Kruskal-Wallis test, and returned a p-value for each of the 10,257 genes considered in our analysis. To control for multiple comparisons and set our false-discovery rate at 0.05, we performed a Benjamini-Hochberg correction on each p-value with alpha set at 0.05 (q-value ≤ 0.05). While this stringent analysis allowed us to infer that we have less than 5% of false positives in our list of 944 genes being misregulated by shift work (944 genes with a q-value ≤ 0.05), it is likely that many other genes are affected by shift work since 2,950 of them have a p-value ≤ 0.05. **A**. Distribution of p-values and q-values of the Kruskal-Wallis test performed between the C, W, and WRF rats for the 10,257 expressed genes in the rat heart. **B**. Number of differentially expressed genes between C, W, and WRF groups based on the p-value threshold (p < 0.01 or p < 0.05) **C**. Description of the significant effects between groups after Mann-Whitney U test post hoc analysis when considering a p-value threshold of 0.05 for the Kruskal-Wallis test. Categories of statistically significant differential gene expression are illustrated by bar graphs with the total number of genes written above for each category. Groups with different letters are significantly different.

**Fig. S4:**
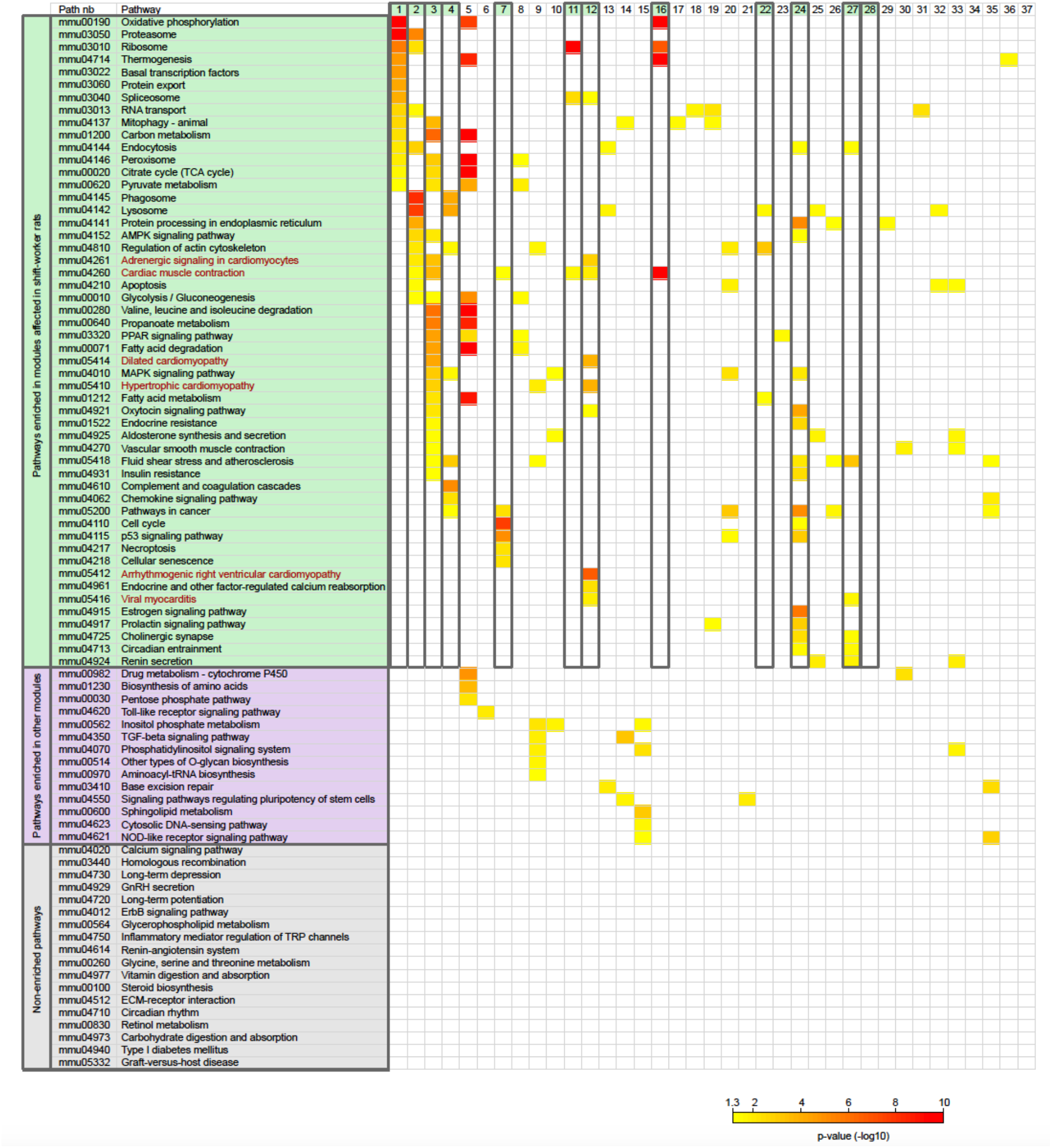
KEGG pathway analysis on modules M0 to M37. Modules from Nomura et al., 2018 correspond to distinct sets of genes being misregulated by cardiac hypertrophy in the mouse heart (24). Genes from each module were retrieved and cross-referenced with our list of 10,257 genes expressed in the rat heart. Genes expressed in the rat heart and categorized within each module were used for a KEGG pathway analysis using the function kegga of the R package Limma. The full list of KEGG pathways is provided in Table S8. Relevant KEGG pathways were selected and their enrichment in modules M1 to M37 are displayed using color-coding. No enrichment is displayed in white.

**Fig. S5:**
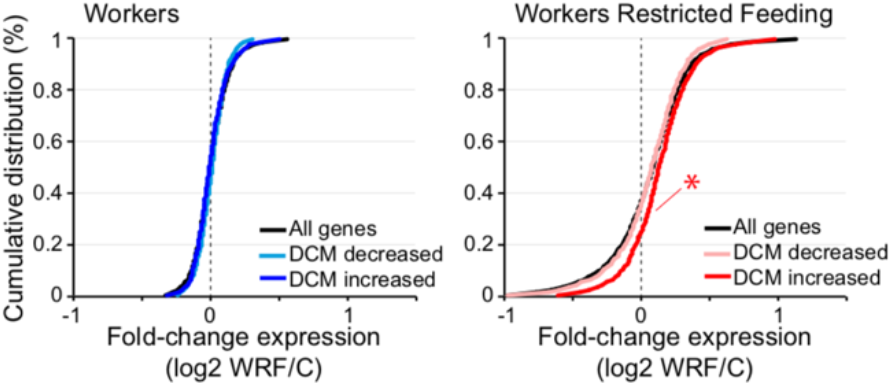
Effect of shift work on the expression of genes misregulated by dilated cardiomyopathy. Cumulative distribution of WRF/control or W/control expression ratio for all genes (black) or for genes described as either downregulated or upregulated by dilated cardiomyopathy in the human heart. The list of down/up-regulated genes was retrieved from Sweet et al., 2018 (37). Distribution significant different from that of all genes is illustrated by an asterisk (Kolmogorov-Smirnov test, p < 0.05).

**Fig. S6:**
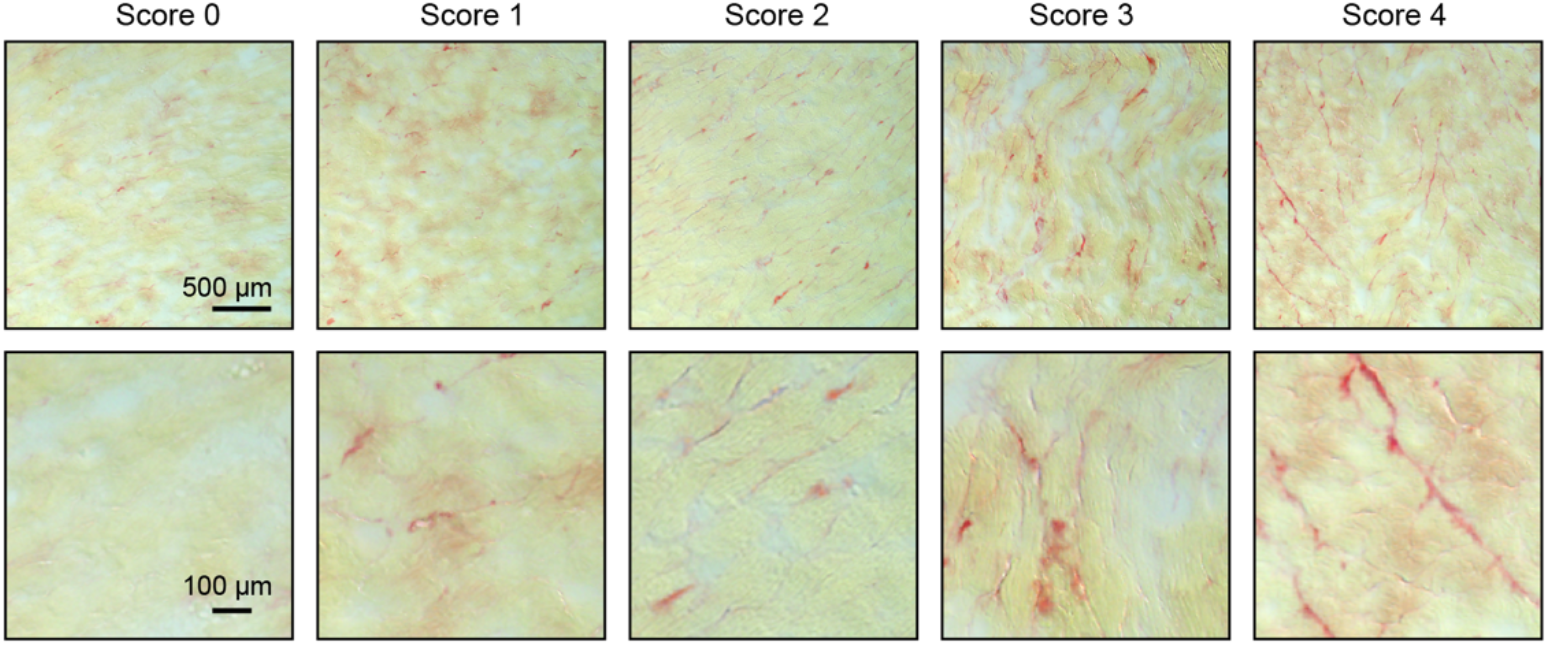
Quantification of picrosirius red staining in the heart. Representative examples of picrosirius red staining in 10 μm sections of rat heart. A score of 0-4 was given for collagen deposition. Yellow =viable myocardium, and red = collagen.

## Notes

https://www.ncbi.nlm.nih.gov/geo/query/acc.cgi?acc=GSE124870

### Competing Interest Statement

The authors have declared no competing interest.

